# Molecular simulations reveal the free energy landscape and transition state of membrane electroporation

**DOI:** 10.1101/2023.01.31.526495

**Authors:** Gari Kasparyan, Jochen S. Hub

## Abstract

The formation of pores over lipid membranes by the application of electric fields, termed membrane electroporation, is widely used in biotechnology and medicine to deliver drugs, vaccines, or genes into living cells. Continuum models for describing the free energy landscape of membrane electroporation have been proposed decades ago, but they have never been tested against spatially detailed atomistic models. Using molecular dynamics (MD) simulations with a recently proposed reaction coordinate, we computed potentials of mean force of pore nucleation and pore expansion in lipid membranes at various transmembrane potentials. Whereas the free energies of pore expansion are compatible with previous continuum models, the experimentally important free energy barrier of pore nucleation is at variance with established models. We trace the discrepancy to previously incorrect assumptions on the geometry of the transition state; previous continuum models assumed the presence of a membrane-spanning defect throughout the process whereas, according to the MD simulations, the transition state of pore nucleation is typically passed before a transmembrane defect has formed. A modified continuum model is presented that qualitatively agrees with the MD simulations. Using kinetics of pore opening together with transition state theory, our free energies of pore nucleation are in excellent agreement with previous experimental data.

## INTRODUCTION

Membrane electroporation is routinely used to form pores in the lipid membrane of biological cells [1–3]. The technique enables the delivery of cargos such as vaccines, drug, genes, or dyes into cells for a wide range of applications in biology, biotechnology, medicine, and food technology [4–6]. The exposure of cells to short electric pulses typically leads to reversible electroporation as the pore may reseal spontaneously [7, 8]. Longer or stronger pulses lead to irreversible electroporation and cell death as used for ablating malignant tumors that are not accessible for surgery [9, 10].

Electroporation has frequently been modeled assuming two distinct lipid arrangements [1, 11, 12]. Accordingly, during pore nucleation, a hydrophobic pore is formed that is characterized by the protrusion of a thin water needle into the membrane core. As the pore radius expands, the lipid headgroups reorient to shield the aqueous defect from the hydrophobic membrane core, thereby forming a hydrophilic pore. Continuum theories for modeling the free energy landscape of this process are considered matured since the late 1980s [7]. The established theory describes the hydrophobic pore as a membrane-spanning cylinder with an unfavorable free energy due to the surface tension at the cylindrical water–membrane interface. The free energy of the hydrophilic pore involves a radius-dependent line tension along the pore rim accounting for the cost for reorienting the lipid headgroups and for the unfavorable lipid packing at the rim [6]. The free energy landscape predicted by this continuum theory has, to the best of our knowledge, never been compared with predictions from spatially more detailed atomistic models.

Molecular dynamics (MD) simulations have been crucial for obtaining atomic insight into membrane electroporation [13–18]. MD simulations confirmed the presence of hydrophobic and hydrophilic pores during pore formation, while the radii of stable open pores were in good agreement with experimental estimates [18]. The effect of lipid composition on pore formation has been investigated in detail by simulations [19–23]. To induce pores within accessible simulation times, pores have been formed under excessive non-equilibrium conditions by applying large transmembrane potentials of several volts, which would lead to membrane rupture after a pore has nucleated. However, because good reaction coordinates (RCs) for driving pore formation were not available until recently, MD simulation did not lead to understanding of the free energy landscape of pore formation. Such understanding would be highly valuable to design electric pulses with desired effects on membranes [24], to rationalize the kinetics of pore opening and closing under conditions of different potentials or different lipid compositions [8, 25], or to translate results found for model membranes into complex biological membranes [23]. Here, we closed this gap and computed the free energy landscape of electroporation covering pore nucleation and expansion at experimentally relevant potentials.

## RESULTS AND DISCUSSION

### Free energy landscape of pore nucleation and expansion

Potentials of mean force (PMFs) were computed along a recently proposed RC *ξ_p_* for pore nucleation and expansion [26]. For *ξ_p_* ⪅ 1, the RC quantifies the degree of connectivity of a polar transmembrane defect (Figs. 1A–C) [27]; for *ξ_p_* ⪆ 1, the RC quantifies the pore radius *R* in units of the radius *R*_0_ ≈ 0.4 nm of a fully nucleated pore, i. e. *ξ*_p_ = *R/R*_0_ (Figs. 1C–E). The transition between nucleation and expansion is implemented with a differentiable switch function (see SI Methods). We showed previously that PMF calculations with this RC do not suffer from hysteresis problems, in contrast to PMF calculations with several other RCs for pore formation [26–28].

**Figure 1:**
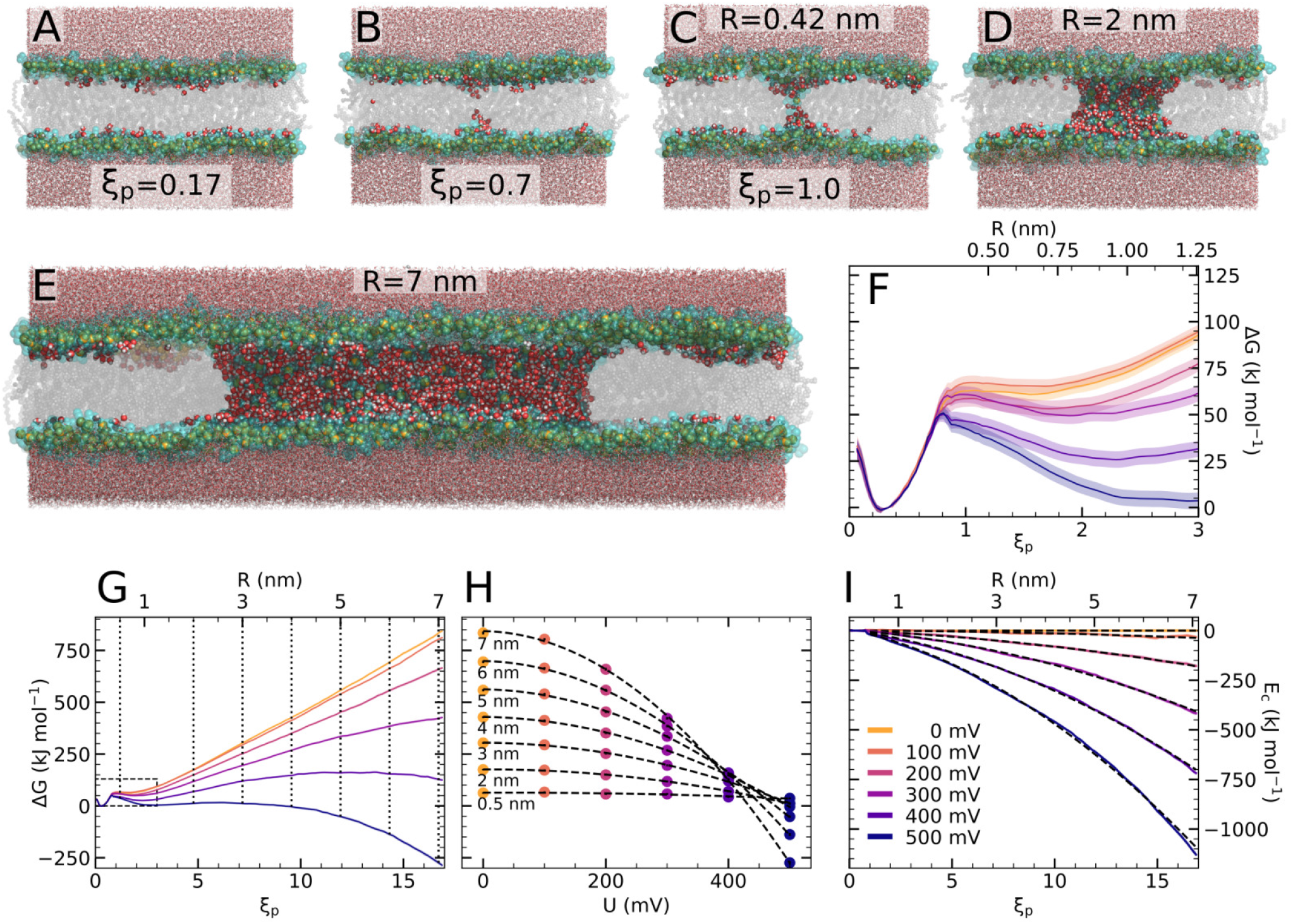
(A–C) Simulation snapshots of pore nucleation and (C–E) pore expansion. Lipid headgroups are shown as green spheres, tails in grey, water within the membrane a red/white spheres, and other water as red/white sticks. The respective position along the reaction coordinate *ξ_p_* and pore radius *R* are shown in the panels. (F/G) Potentials of mean force (PMFs) for pore nucleation and pore expansion across a DPPC membrane at transmembrane voltages between 0 and 500 mV (for color code see panel I). Lower and upper abscissas show the reaction coordinate and the pore radius, respectively. The dashed box in panel (G) indicates the region highlighted in panel (F). (H) Pore free energy versus voltage for pore radii between 0.5 and 7 nm (see labels and vertical lines in panel G). Dashed lines show quadratic fits. (I) Electrostatic stabilization of the pore relative to the PMF of 0 mV. Dashed lines show quadratic fits.

Figures 1F/G present PMFs for electroporation across a membrane of dipalmitoylphosphatidylcholine (DPPC) at transmembrane potentials *U* between 0 and 500 mV, modeled with the force field by Berger *et al*. [29]. The PMFs reveal a free energy cost of 50 to 65 kJ/mol for the formation of a thin water needle as required for pore nucleation. The presence of free energy minima around *R* ∼ 0.8 nm demonstrate the presence of metastable, long-living pores for voltages *U* ≤ 500 mV, while this free energy minimum is very shallow at zero voltage (Fig. 1F, yellow). With increasing voltage, the minimum becomes more pronounced and is shifted to larger *R*, indicating that the metastable pores exhibit increasing lifetime and conductivity. At *U* ≥ 500 mV, the PMFs exhibit no significant barrier against further expansion (Figs. 1F/G, dark blue), suggesting that the pores spontaneously further expand leading to membrane rupture. Free simulations confirmed that pores are metastable at intermediate voltages, whereas pores spontaneously close or expand at small or large voltages, respectively, in agreement with the PMFs (Supplementary Information, Figs. S2 and S3).

Continuum theories of pore expansion suggest that the transmembrane potential modifies the pore free energy by the electrostatic energy *E*_c_ = −Δ*CU* ^2^/2, where the change of the capacitance of the membrane is Δ*C* = (*∊_w_ − ∊_m_*)*πR*^2^/*d*. Here, *∊_w_* and *∊_m_* denote the dielectric permittivity of water and of the membrane core, and *d* is the thickness of the membrane. For the regime of pore expansion (*ξ_p_ >* 1), the PMFs are in agreement with the continuum model of pore expansion as the free energies for fixed pore radii decrease with *U* ^2^ (Fig. 1H), while the PMFs at various voltages are decreased relative to the PMF at zero voltage following mainly a quadratic *R*^2^ dependence (Fig. 1I). Hence, for fully established nanometer-sized pores, the continuum model captures the trends obtained from the MD simulations.

During pore nucleation, however, polar defects with a diameter of few Ångströms protrudes the membrane, suggesting that the finite size of water molecules becomes increasingly relevant. To reveal how the free energy barrier and the transition state (TS) for pore nucleation depend on the voltage, we computed PMFs of pore nucleation across a DPPC membrane using voltages between 0 mV and 1900 mV (Fig. 2A). The PMFs were computed along the chain reaction coordinate *ξ*_ch_, which is a measure for the degree of connectivity of the defect and which is equivalent to *ξ_p_* for *ξ_p_* < 0.9 (SI Methods). Here, *ξ*_ch_ ≈ 0.3 corresponds to the flat unperturbed membrane (Fig. 1A), whereas *ξ*_ch_ ≈ 0.9 indicates a membrane-spanning polar defect (Fig. 2D); pore expansion is not resolved by *ξ*_ch_ since larger pores are projected onto *ξ*_ch_ = 1.

**Figure 2:**
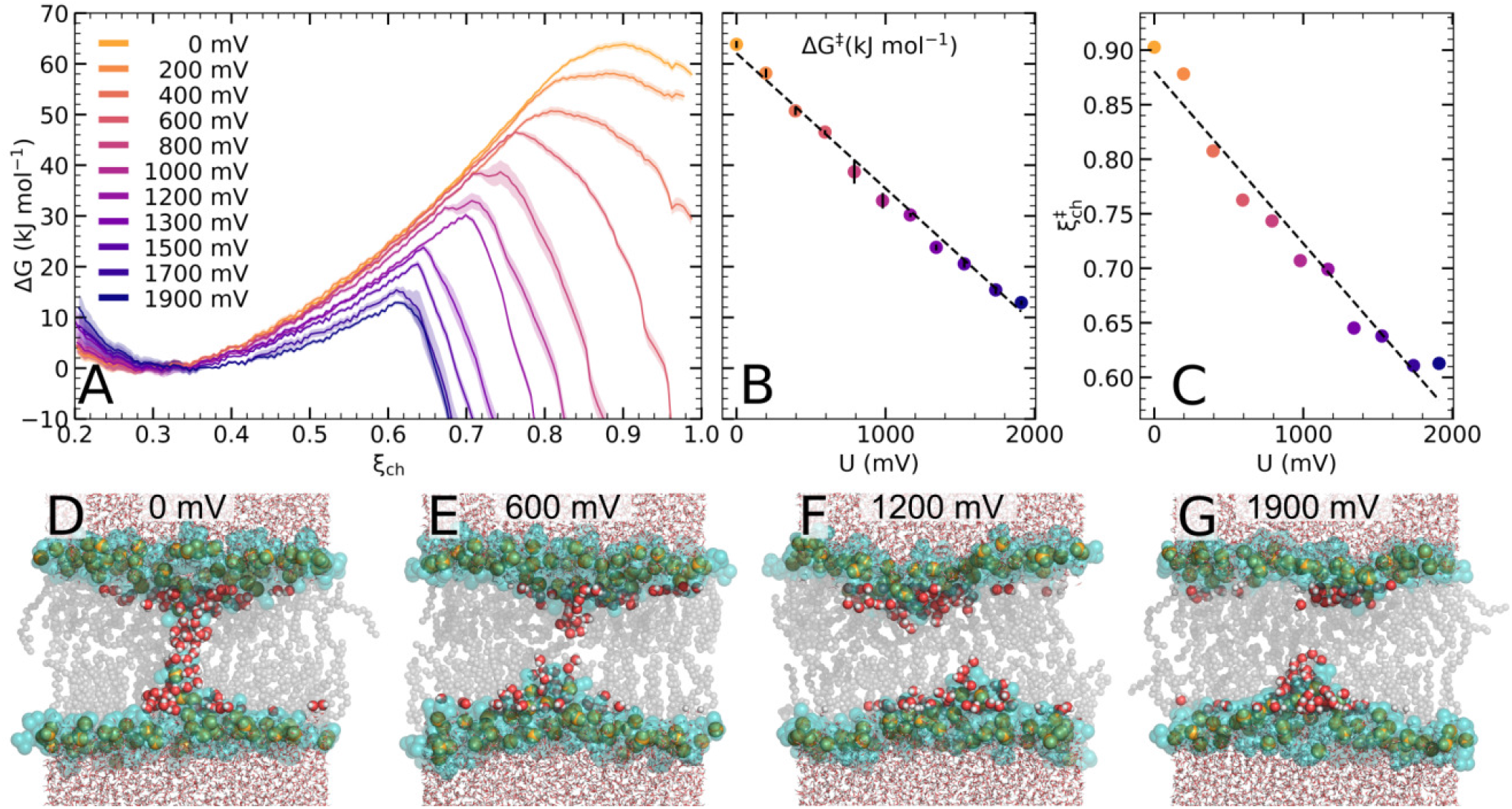
(A) PMFs of pore nucleation at transmembrane voltages between 0 and 1900 mV (see legend) as function of the chain coordinate *ξ*_ch_, which quantifies the connectivity of the transmembrane defect. (B) Free energy barrier for pore nucleation as taken from the maxima of the PMFs in panel (A). (C) Position of *ξ*_ch_ at the transition state. Dashed lines in panels (B) and (C) are linear fits to guide the eye. (D–G) MD snapshots of the transition state, revealing structures with a discontinuous, partial defects.

The PMFs in Fig. 2A demonstrates that the free energy landscape of pore nucleation is greatly modulated by transmembrane voltages. With increasing *U*, the barrier of the PMFs, corresponding to the TS of pore nucleation, is shifted to smaller free energies and to smaller values of *ξ*_ch_. Specifically, the barrier height Δ*G^‡^* of the TS decrease approximately linearly with *U* (Fig. 2B). Hence, not only the open pore is stabilized by the potential, but the kinetics of pore formation are greatly accelerated. Assuming transition state theory (TST), the rate of pore nucleation follows exp(−*β*Δ*G^‡^*), where *β* is the inverse temperature; according to TST, the decrease of the barrier from 64 kJ/mol to 13 kJ/mol upon the application of 1900 mV implies that the rate of pore formation is accelerated by approximately eight orders of magnitude. Notably, the linear decay of Δ*G^‡^* with *U* is at variance with widely accepted continuum models discussed below [6, 11, 12, 25, 30, 31]; however, the linear decay has been proposed previously by Böckmann *et al*. by extrapolating kinetics of pore formation observed in MD simulation at large potentials towards experimental kinetics at small potentials [18].

The position of the TS along the reaction coordinate, 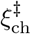, is shifted to smaller values with increasing voltage (Fig. 2C). Since *ξ*_ch_ quantified the degree of connectivity of the defect, this implies that, with increasing potential, the TS is characterized by a decreasingly connected polar defect. Whereas the TS for 0 mV represents a continuous transmembrane defect (Fig. 2D), the TSs for higher potentials exhibit only a partial water protrusions that span the membrane incompletely (Figs. 2E–G). Hence, during pore nucleation at higher potentials, the TS is passed *before* a continuous transmembrane needle has formed. This finding suggests that previous continuum models, which modeled the entire pore formation process using transmembrane cylinders with increasing radii, were not capable of modeling the TS at higher potentials.

### Revised continuum model of pore nucleation

Previous continuum theories modeled pore formation via a transition from a hydrophobic to a hydrophilic membrane-spanning defect (Figs. 3A/B) [1, 7, 11, 32], as reviewed repeatedly [6, 25, 30, 31]. The free energy of the hydrophobic defect was taken as Δ*G*_o_(*R, U*) = 2*πRd*Γ_eff_ (*R*) + *E*_c_, where Γ_eff_ denotes an effective radius-dependent surface tension between a membrane-spanning polar cylinder of radius *R* and height *d* in contact with the hydrophobic membrane core (Fig. 3B, left inset). For larger pores, the lipid headgroups reorient to shield the polar defect from the membrane core, thereby forming a hydrophilic pore with free energy Δ*G*_i_(*R, U*) = 2*πRγ*(*R*) + *E*_c_. Here, *γ*(*R*) is the line tension along the pore rim accounting for the cost for tilting the lipids and for remaining water–tail interactions; the line tension was empirically taken as *γ*(*R*) = *γ*(1+*C/R*^5^) with parameters *γ* and *C* to model the increasingly unfavorable curvatures of small hydrophilic pores [6]. Δ*G*_i_(*R, U*) may, in addition, contain an energy term for a release of surface tension; such term is not considered in this study since we (i) simulated without external tension and (ii) at constant number of lipids, suggesting that the surface tension contribution owing to the membrane–water interface remains approximately constant. According to this theory, the overall free energy profile would be given by the smaller value among Δ*G*_o_(*R, U*) and Δ*G*_i_(*R, U*), as shown in Fig. 3A for voltages between 0 and 500ṁV, using the parameters in Table SI. The TS of pore formation would be located at the radius *R*_min_, where Δ*G*_o_ = Δ*G*_i_, indicating the radius of the transition from the hydrophilic to the hydrophobic pore, and visible in the profiles of Figs. 3A/B as a marked maximum at *R*_min_ ≈ 0.5 nm. The free energy barrier would decay quadratically with increasing potential (Fig. 3C), while the position *R*_min_ and, thereby the structure of the TS, is independent of *U*. These properties are at variance with our MD simulation that suggest an approximately linear decay of the TS free energy (Fig. 2B) and a potential-dependent TS structure (Figs. 2C–G).

**Figure 3:**
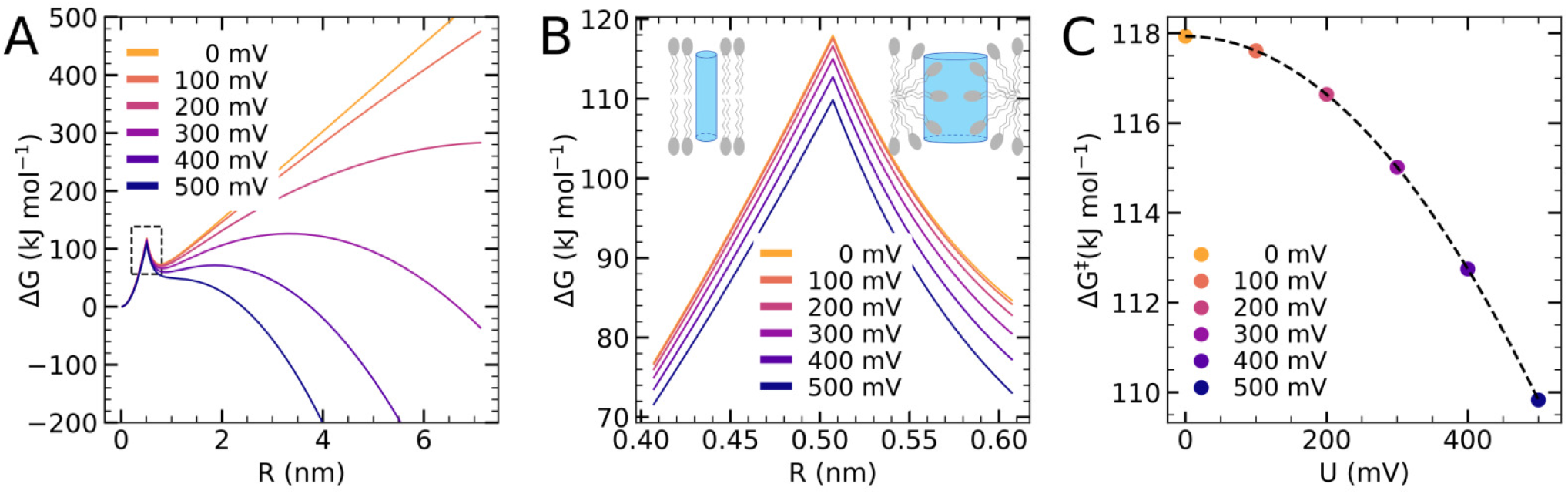
Free energy landscape of electroporation according to previous continuum models assuming a transition from a hydrophobic to a hydrophilic membrane-spanning cylindrical defect. PMFs are shown for potentials between 0 mV and 500 mV, at zero surface tension. (A) Overall PMF versus pore radius, (B) closeup view on the transition region (cf. small dashed box in panel A). The left branches of the profiles model assume a hydrophobic defect (*R* ≲ 0.5 nm; left inset in panel B); the right branches assume a hydrophilic defect (*R* ≳ 0.5 nm; right inset in panel B) (C) Quadratic decrease of the free energy barrier.

The discrepancy between MD simulation and previous continuum models is explained by different molecular conformations of the TS. Whereas previous continuum models assumed membrane-spanning water cylinders throughout the pore formation process, MD simulations suggest that, at higher potentials, the TS is passed before a continuous defect has formed. Hence, we devised a modified continuum model involving a gradual formation of a transmembrane defect and, thereby, capable of capturing the TS of pore nucleation. We model the growing defect as three capacitors in series, where the upper and lower cylinder-shaped capacitors of height *h*/2 are filled by water, whereas the central capacitor is filled by lipid tails (Figs. 4 and 6). As detailed in Appendix I, the free energy change upon protrusion of the two water cylinders is

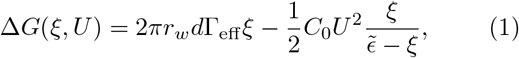

where *r_w_* is the constant radius of the cylinder, *C*_0_ the capacitance of a purely hydrophobic cylinder, 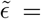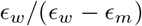, and *ξ* = *h/d* ∈ [0, 1] the degree of connectivity of the defect, similar to the chain coordinate *ξ*_ch_ used in the PMF calculations of pore nucleation. Figure 4B presents Δ*G*(*ξ, U*) curves for potentials between 0 and 4500 mV, revealing a decreasing TS free energy Δ*G^‡^* and a decreasing position *ξ^‡^* with increasing potential, in qualitative agreement with the PMFs from MD simulations (Figs. 2A–C). The transition state given by the maxima of the Δ*G*(*ξ, U*) curves, is at

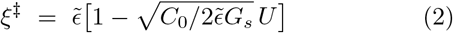

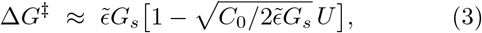

where *G_s_* = 2*πr_w_d*Γ_eff_ denotes the surface tension free energy of a membrane-spanning water-filled cylinder. Hence, *ξ^‡^* and Δ*G^‡^* decrease linearly with the potential (Figs. 4B/C), in agreement with the findings from the PMFs (Figs. 2B/C), suggesting that the model accounts for the correct geometry of pore nucleation and for the dominating energetic contributions.

**Figure 4:**
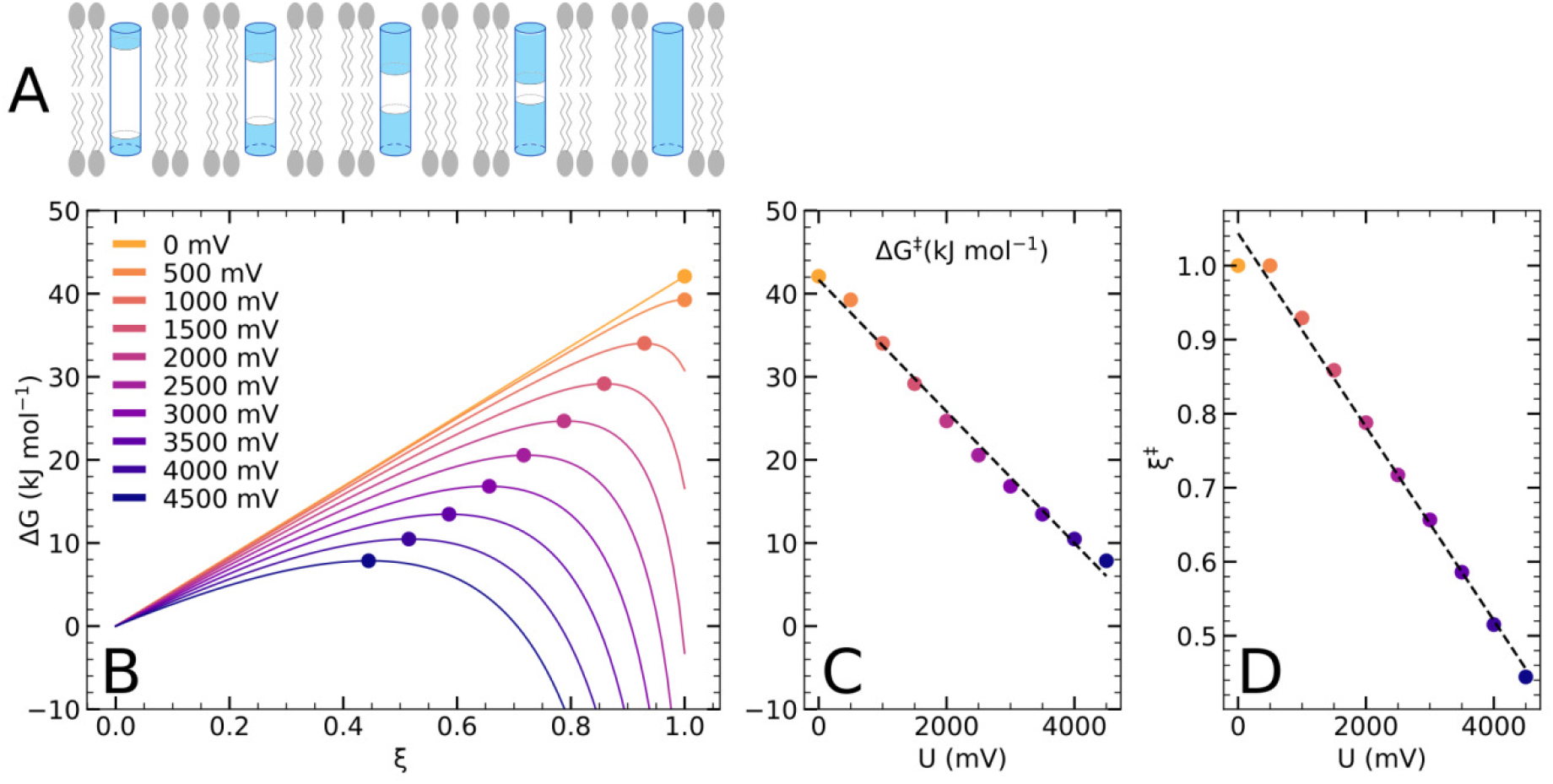
Energetics of pore nucleation by the three-cylinder continuum model. (A) Gradual formation of a transmembrane defect modeled by three cylindrical capacitors in series. (B) Free energy of pore nucleation for transmembrane voltages between 0 and 4500 mV (for color code, see legend); (B) free energy barrier and (C) position of the transition state (colored dots in panel B).

Nevertheless, the thee-cylinder model is simplified as it neglects several contributions to the free energies. First, local elastic thinning of the membrane may stabilize the early water protrusions, which may explain the quadratic free energy increase in the PMFs (Fig. 2A, *ξ*_ch_ ≈ 0.3) in contrast to the linear increase proposed by the three-cylinder model (Fig. 4B, *ξ* ≈ 0). Second, during the MD simulations, individual lipid head groups frequently followed the water protrusions far into the membrane core, even before continuous transmembrane defect has formed; owing to the large permittivity of the head group region [33], such headgroups may increase the overall permittivity of the defect, thereby increasing the electrostatic stabilization *E_c_* of the pore at the cost of some lipid deformation energy. Such effects may rationalize why Δ*G^‡^* decays more rapidly with *U* in the simulations as compared to the three cylinder model (Fig. 2B and Fig. 4C). Early occasional protrusions of individual head groups further explain why the MD simulation do not reveal an instantaneous transition from a hydrophobic to a hydrophilic pore, as anticipated by the previous continuum model (Fig. 3B), but instead a gradual transition from a water needle stabilized by local membrane thinning towards a fully established toroidal pore, as illustrated in Supplementary Movies S1–S6.

### Transition state theory of pore formation

Since the PMFs of pore nucleation exhibit a single barrier (Figs. 1F and 2A), we used TST to describe the rate of pore opening by *k*_o_ = *κ* exp(*β*Δ*G^‡^*), where *κ* is the attempt frequency. We estimated the attempt frequency *κ* from a series of free MD simulations starting with a planar membrane. Within 20 1-μs-simulations each at potentials of 1500 mV, 1700 mV, or 1900 mV, we observed 1, 9, or 19 opening events within simulation time, respectively. Using a maximum-likelihood estimate, we obtained the rates *k*_o_ of pore opening for these voltages (SI Methods, Tab. SII) and, together with the respective free energy barriers (Fig. 2A), the attempt frequency via *κ* = *k*_o_ exp(+*β*Δ*G^‡^*). The attempt frequencies estimated from simulations at different voltages were in reasonable agreement (Tab. SII) suggesting that TST is applicable and yielding an attempt frequency of *κ* ≈ 0.25 ns^*−*1^, corresponding to roughly one attempt per 4 ns.

Notably, the attempt frequency of 0.25 ns^*−*1^ is approximately 3 orders or magnitude smaller as compared to a previous estimate based on the number of lipid collisions [34]. Instead, the time of ~4 ns per attempt corresponds approximately to the time required for lipids to diffuse a typical lipid–lipid distance and, therefore, may be taken as a time scale of lipid–lipid rearrangements. A similar attempt frequency of ∼0.3 ns^*−*1^ was previously derived for describing the kinetics of stalk formation with TST [35]. This suggests that lipid–lipid re-arrangements, rather than lipid collisions, may be interpreted as attempts of large-scale topological transitions of membranes such as pore formation or stalk formation in the context of TST.

### Validation against experimental data

Having established the attempt frequency of pore formation, TST enables estimates of pore opening rates for voltages that do not trigger spontaneous pore opening in free simulations within acceptable simulations times. Following such strategy, we validated the MD simulations of pore formation against experimental data by Melikov *et al*. [36], who reported mean times of pore opening across black lipid membranes of diphytanoylphosphocholine (DPhPC) at voltages between 250 mV and 550 mV. To this end, we computed PMFs of pore nucleation across a membrane of DPhPC at voltages between 0 mV and 1200 mV using the Charmm36 force field (Fig. 5A, SI Methods) [37]. From the free energy of the pore (*ξ*_ch_ ≈ 0.9) or from the free energy barrier (if present) together with TST, we computed the mean time of pore opening *τ*_o_ for a circular black lipid membrane with radius of 150 μm [36]. The mean pore opening times *τ*_o_ from the MD simulations are in good agreement with experimental data both in terms of the magnitude and in terms of the voltage-dependence of *τ*_o_ (Fig. 5B). The agreement is remarkable considering that neither the applied Charmm36 force field has been refined against free energies of pore opening nor any other fitting has been applied for this comparison between simulation and experiment.

**Figure 5:**
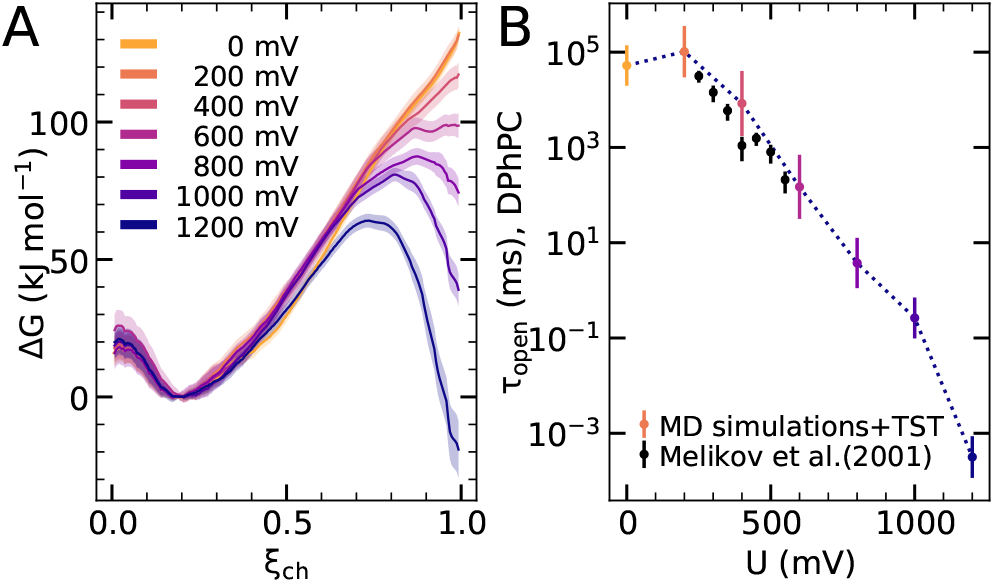
(A) PMFs of pore nucleation in a DPhPC membrane at voltages between 0 mV and 1200 mV (see legend). (B) Mean time of pore opening derived from MD simulations based on transition state theory and using the pore free energies (or free energy barriers, if present) in panel A (colored dots). Black dots: Experimental data by Melikov *et al*. [8].

**Figure 6:**
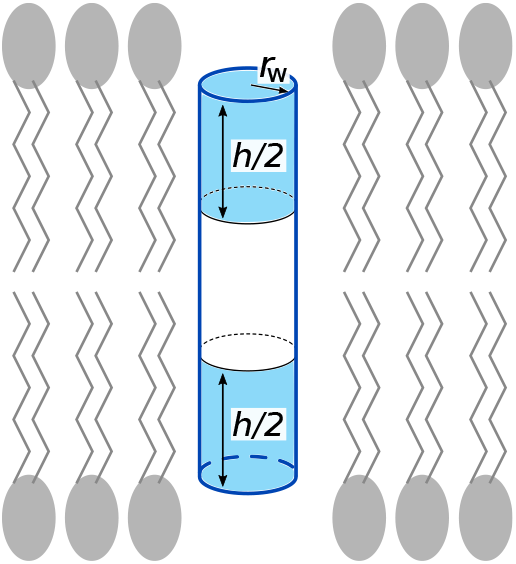
Pore nucleation model of electroporation composed of three cylinders. The blue cylinder segments indicate water-filled regions with dielectric permittivity *ϵ_w_*, while the white segment indicates a hydrophobic region with dielectric permittivity *ϵ_m_*.

## CONCLUSIONS

To conclude, we derived the free energy landscape of membrane electroporation from all-atom MD simulations, involving both pore nucleation and pore expansion. We showed that previous continuum theories, which modeled pore formation purely based on a continuous transmembrane water cylinder, are in good agreement with the MD-based PMFs only in the pore expansion regime, that is, after a continuous transmembrane defect has formed. However, previous models failed to explain the free energy and structure of the TS of pore nucleation because, according to the MD simulations at higher potentials, the TS is passed before a membrane-spanning defect has formed. We presented a continuum model of pore nucleation that agrees qualitatively with the MD-based PMFs of pore nucleation. Finally, using TST together with (i) an attempt frequency corresponding to the time scale of lipid–lipid rearrangements and free energies from the PMFs, we derived pore opening times in quantitative agreement with previous experimental data. The comparison of pore formation kinetics between simulation and experiments validates not only the PMF calculation protocol, but furthermore provides a new means for testing the accuracy of lipid force fields under large-scale membrane conformational transitions. Together, this study provided energetic and structural insight into membrane electroporation and paves the way for interpreting and designing electroporation applications in biotechnology and medicine.

## APPENDIX I: THREE-CYLINDER CONTINUUM MODEL OF PORE NUCLEATION

## Free energy versus connectivity of the defect

We model pore nucleation by the formation of two cylinder-shaped water needles that penetrate the hydrophobic membrane core symmetrically from the upper and the lower water compartments (Fig. 6). For the sake of simplicity, local membrane thinning owing to partial bending of the headgroups towards the membrane core is neglected. Let *d* denote the thickness of the membrane core and *h*/2 the depth of the two penetrating water needles. The coordinate *ξ* = *h/d* ∈ [0, 1] is the fraction to which the membrane is penetrated and, thereby, quantifies the degree of connectivity of the transmembrane defect, qualitatively similar to the chain coordinate *ξ*_ch_ applied in our MD simulations of pore nucleation.

Hence, the nucleation site may be modeled by three capacitors in series with thicknesses *h*/2, *d − h*, and *h*/2, respectively. The capacitance of the two cylinder-shaped water needles equals *C_w_* = *ϵ_w_A*/(*h*/2), and the capacitance of the central hydrophobic cylinder is *C_m_* = *ϵ_m_A*/(*d − h*), where *ϵ_w_* and *ϵ_m_* denote the dielectric permittivities of water and of the membrane core, respectively. 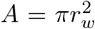 is the cross section area of the cylinder with radius *r_w_*.

The total capacitance of the nucleation site is

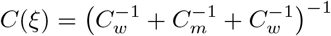

The change of capacitance Δ*C*(*ξ*) of the nucleation site relative to the capacitance *C*_0_ ≔ *C*(0) = *ϵ_m_A/d* of the flat unperturbed membrane is

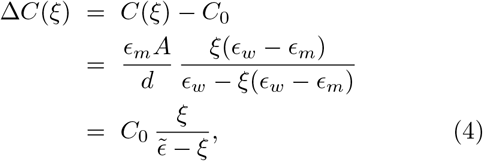

where we introduced 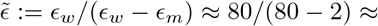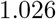. For the states with the unperturbed membrane (*ξ* = 0) and with a continuous water needle (*ξ* = 1), Δ*C*(*ξ*) in Eq. 4 simplifies as expected to Δ*C*(0) = 0 and Δ*C*(1) = (*ϵ_w_ ϵ_m_*)*A/d*, respectively.

As the two water needles penetrate the membrane and, thereby, nucleate the pore, the free energy is modified by two contributions. First, because the lipids are still oriented upright (Fig. 6), the penetrating water is in contact with the hydrophobic membrane core, leading to unfavourable interface. The free energy is increased by

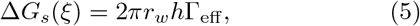

where Γ_eff_ is the effective surface tension between water needle and membrane core, which differs from the surface tension Γ_*∞*_ of a macroscopic interface between water and hydrophobic core owing to the large curvature of the cylinder surface. Following previous work [6, 38], we take Γ_eff_ = Γ_*∞*_*I*_1_(*r_w_/λ*)/*I*_0_(*r_w_/λ*), where *I_k_* denotes Bessel functions of *k*^th^ order, and *λ* ≈ 1 nm is the characteristic length of hydrophobic interactions. Note that the contributions at the two cylinder faces (with areas *A*) are not considered here since these faces was already present in the initial state at *ξ* = 0.

Second, the electrostatic energy decreases by

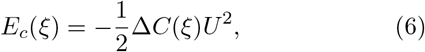

where *U* is the transmembrane potential. Here, the minus sign corresponds to the observation that a dielectric medium is pulled into a capacitor given that the capacitor is kept at constant voltage by a power source. According to Eqs. 4–6, the total free energy as function of pore nucleation *ξ* is

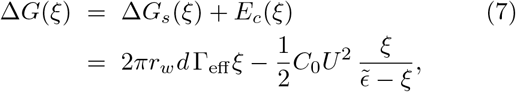

where we used *h* = *ξd*. Typical curves for Δ*G*(*ξ*) for transmembrane potentials between 0 and 4.5 V in steps of 0.5 V are shown in Fig. 4A, using the parameters in Table SI.

## Transition state free energy and position

Critically, the Δ*G*(*ξ*) curves reveal maxima corresponding to the transition states of pore nucleation, and these maxima are located at *ξ* < 1. Hence, at the transition state, the two penetrating water needles are not yet connected but instead only partly penetrate the membrane. In other words, the transition state is passed *before* a continuous transmembrane defect has formed. This suggests that previous models of electroporation based purely on continuous membrane-spanning cylinders (Fig. 3B) have not been capable of modeling the transition state of pore nucleation.

To find the maximum of Δ*G*(*ξ*) corresponding to the nucleation free energy Δ*G^‡^*, we rewrite Eq. 7 as

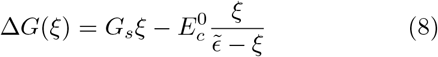

with abbreviations *G_s_* ≔ 2*πr_w_d* Γ_eff_ and 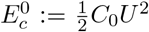. Solving d(Δ*G*)/d*ξ* = 0 yields the maximum of Δ*G*(*ξ*) at

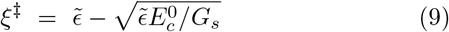

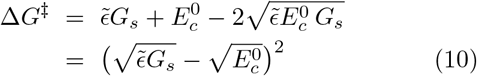

TSs defined by (*ξ^‡^*, Δ*G^‡^*) are illustrated in Fig. 4A as circles. Figs. 4B and C present Δ*G^‡^* and *ξ^‡^* as function of potential *U*, respectively, revealing approximately linear decays in the relevant potential range, in qualitative agreement with the PMFs obtained from our MD simulations (Figs. 2B/C).

It is instructive to rewrite Eqs. 9 and 10 as functions of transmembrane potential *U* and using that, for smaller or moderate potentials, 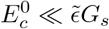. We obtain

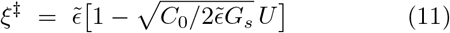

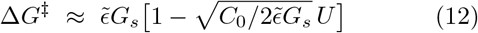

demonstrating that both *ξ^‡^* and Δ*G^‡^* decrease (approximately) linearly with *U*. Hence, according to Eq. 11, the larger the potential, the earlier (at lower *ξ*) the transition state is reached. Noteworthy, a barrier is only present if *ξ^‡^* < 1, that is, only if the transmembrane potential *U* is above the critical value

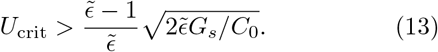

## Acknowledgements

This study was supported by Deutsche Forschungsgemeinschaft (DFG, German Research Foundation; grants SFB 803/A12 and SFB 1027/B7).

