## Supplemental Information for "Molecular simulations reveal the free energy landscape and transition state of membrane electroporation"

Gari Kasparyan[1] and Jochen S. Hub[1]*
[1] *Theoretical Physics and Center for Biophysics, Saarland University, 66123 Saarbrücken, Germany*

### I. KINETICS OF PORE OPENING FROM FREE SIMULATIONS

To obtain the kinetics of pore opening, 20 simulations of 1 µs each were carried out with transmembrane potentials 1500 mV, 1700 mV, or 1900 mV, using the 128 DPPC lipid simulation system (see below for details). Simulations were started from the final simulation frame of an umbrella sampling simulation with $U = 1700\,\mathrm{mV}$ restrained to $\xi_{\mathrm{ch}} = 0.24$. Hence, the initial conformation was pre-equilibrated in the presence of a larger transmembrane potential.

Among the 20 1 µs simulations, we observed 1, 9, and 19 opening events in the simulations with 1500 mV, 1700 mV, and 1900 mV, respectively (Tab. SII). Assuming that the transition from the planar membrane to the pore follows a simple monoexponential decay, we derived the maximum-likelihood estimate of the lifetime $\tau_\mathrm{o}$ until pore opening. Similar to previous work [S2], $\tau_\mathrm{o}$ was estimated from the number and the simulation times of the opening events:

$$\tau_\mathrm{o} = \frac{1}{n_\mathrm{o}}\left[(N_\mathrm{sim} - n_\mathrm{o})T_s + \sum\nolimits_{i=1}^{n_\mathrm{o}} t_{\mathrm{o},i}\right] \qquad \text{(S1)}$$

Here, $N_\mathrm{sim} = 20$ is the number of simulations, $n_\mathrm{o}$ the number of opening events, $T_s = 1\,\mu\mathrm{s}$ the length of the simulations, and $t_{\mathrm{o},i}$ are the times at which pore opening occurred. The rate of pore opening is given by $k_\mathrm{o} = 1/\tau_\mathrm{o}$. Assuming transition state theory, $k_\mathrm{o} = \kappa \exp(-\beta\Delta G^\ddagger)$, we estimated the attempt frequencies $\kappa$ using the barriers $\Delta G^\ddagger$ from the PMFs together with the opening rates $k_\mathrm{o}$ from the free simulations.

The results, as summarize in Table SII, suggest attempt frequencies in the order of $0.25\,\mu\mathrm{s}^{-1}$, corresponding to approximately one attempt per 4 ns. Here, the attempt frequencies derived from the 1700 mV and 1900 mV simulations are statistically more robust as compared to the estimate from the 1500 mV simulations, since the former are based on 9 and 19 opening events, respectively, whereas the latter is based on only a single opening event. Nevertheless, the attempt frequencies $\kappa$ obtained from different potentials show reasonable agreement, considering that the computed values of $\kappa$ are sensitive with respect to uncertainties of $\Delta G^\ddagger$.

Having established the attempt frequency $\kappa$, and to validate our simulations against experimental data by Melikov *et al.* [S3], we used TST to predict the mean time for pore opening $\tau_\mathrm{o}$ at lower potentials for a macroscopic membrane with area $A_\mathrm{exp}$. Using that the probability of pore opening is proportional to the membrane area, we used $\tau_\mathrm{o}^{-1} = \kappa A_\mathrm{exp}/A_\mathrm{sim} \times \exp(-\beta\Delta G^\ddagger)$. To obtain the results shown in Fig. 5B, we assumed a circular experimental membrane with a radius 150 µm [S3] and used for the membrane area in the simulations $A_\mathrm{sim} = (6.6\,\mathrm{nm})^2$.

Table SI: Numerical values used for three-cylinder model of pore nucleation proposed here and for the previous continuum models

| Symbol | value | Ref. |
|---|---|---|
| $d$ | 5 nm | - |
| $r_w$ | 0.3 nm | - |
| $\Gamma_\infty$ | $5 \times 10^2\,\mathrm{J/m^2}$ | [S1] |
| $\lambda$ | 1 nm | [S1] |
| $\epsilon_w$; $\epsilon_m$ | 80; 2 | - |
| $\gamma$ | $1 \times 10^{-11}\,\mathrm{Jm}$ | [S1] |
| $C$ | $1.39 \times 10^{-46}\,\mathrm{Jm^4}$ | [S1] |

### II. COMPUTATIONAL DETAILS

#### A. Simulation setup and parameters

Simulation systems were prepared using MemGen [S4]. Three systems with DPPC lipids were simulated. For pore nucleation, we used a small system of 128 lipids and 40 water molecules per lipid (21,760 atoms, Figs. 2D–G). For pore expansion, a medium-sized system with 512 lipids and 60 water molecules per lipid was used (117,760 atoms, Figs. 1A–D) as well as a large system with 2048 lipids and 60 water molecules per lipid (471,040 particles, Fig. 1E). DPPC lipids were described with the force field by Berger *et al.* [S5] and the SPC water model was used [S6]. The systems were equilibrated with the Gromacs simulation software, versions 2020.1 or 2021.5 [S7]. The integration time step was set to 4 or 5 fs. The temperature was controlled at 323 K by velocity rescaling using water and lipids as separate coupling groups ($\tau = 0.1\,\mathrm{ps}$) [S8]. The pressure was controlled at 1 bar

*Electronic address:

Table SII: Kinetics of pore opening from free simulations. Transmembrane potential $U$, number of simulations $N_{\mathrm{sim}}$ of 1 µs each, number of opening events within 1 µs of simulation time $n_{\mathrm{o}}$, average simulation time at pore opening events $t_{\mathrm{o}}^{\mathrm{av}}$, maximum-likelihood estimate of opening rate $k_{\mathrm{o}}$, barrier for pore opening $\Delta G^{\ddagger}$ taken from the PMFs, attempt frequency of pore opening $\kappa$ in the context of transition state theory, $k_{\mathrm{o}} = \kappa \exp(-\beta \Delta G^{\ddagger})$

| U(mV) | $N_{\mathrm{sim}}$ | $n_{\mathrm{o}}$ | $t_{\mathrm{o}}^{\mathrm{av}}$ (ns) | $k_{\mathrm{o}}$ (µs$^{-1}$) | $\Delta G^{\ddagger}$ (kJ/mol) | $\kappa$ (1/ns) |
|---|---|---|---|---|---|---|
| 1500 | 20 | 1 | 450 | 0.05 | 20.7 | 0.2 |
| 1700 | 20 | 9 | 404 | 0.61 | 15.5 | 0.31 |
| 1900 | 20 | 19 | 310 | 3.2 | 12.5 | 0.21 |

with a semi-isotropic Berendsen barostat ($\tau = 1$ ps) during pore nucleation and, at later stage of this project, using cell rescaling ($\tau = 5$ ps) during pore expansion [S9]. Lennard-Jones interactions were truncated at 1 nm. Electrostatic interactions were calculated using the particle-mesh Ewald method [S10, S11]. Water molecule bonds and angles were constrained with SETTLE [S12] and lipid bonds were constrained with LINCS [S13]. The systems were equilibrated for at least 40 ns.

Transmembrane potentials were generated by applying an external electric field $E_z = U/L_z$ perpendicular to the membrane plane, $L_z$ is the box length in $z$ direction (membrane normal) [S14, S15]. We recently showed that using either (i) external fields or (ii) charge imbalance between the solvent reservoirs of a stacked double-membrane system leads to identical PMFs of membrane electroporation and identical water dipole orientations within transmembrane defects [S16]. Hence, using an external field is a valid approach for modeling transmembrane potentials during electroporation.

During the simulation of the DPhPC membrane, the lipids were described by the Charmm36 force field [S17, S18] and using hydrogen mass repartitioning [S19]. Water was modeled with the Charmm-modified TIP3P model [S17, S20]. Lennard-Jones interactions were treated with a force switch between 1 and 1.2 nm. A time step of 4 fs was used. The temperature was controlled at 310 K. Other parameters were chosen as described above.

#### B. Reaction coordinates for pore nucleation and pore expansion

PMFs were computed using umbrella sampling (US) [S21] along tailor-made reaction coordinates (or collective variables) for pore nucleation and pore expansion. PMFs of pore nucleation (Fig. 2A) were computed along the chain coordinate $\xi_{\mathrm{ch}}(\mathbf{r})$ introduced previously [S2, S22], where $\mathbf{r}$ denotes the Cartesian coordinates of the simulation atoms. PMF calculations along $\xi_{\mathrm{ch}}$ have been used to quantify the effects of membrane-active peptides, polymers, or drugs on the free energy of pore formation [S23–S25]. The coordinate $\xi_{\mathrm{ch}} \in [0, 1)$ quantifies the connectivity of a polar defect that penetrates the membrane hydrophobic core. $\xi_{\mathrm{ch}}$ is defined with the help of a membrane-spanning cylinder that is decomposed in to $N_s$ slices. The coordinate has been defined as

$$\xi_{\mathrm{ch}}(\mathbf{r}) = \frac{1}{N_s} \sum_{s=0}^{N_s-1} \delta_\zeta(n_s^{(p)}), \tag{S2}$$

where $n_s^{(p)}$ is the number of polar atoms in cylinder slice $s$. $\delta_\zeta \in [0, 1)$ is a differentiable approximation to an indicator function, which takes 0 for an empty slice ($n_s^{(\mathrm{p})} = 0$) and a value close to unity for a slice filled by polar atoms ($n_s^{(p)} \geq 1$).To render $\xi_{\mathrm{ch}}$ differentiable with respect to the Cartesian coordinates $\mathbf{r}$, $n_s^{(p)}$ is defined using differentiable switch functions. For more details on $\xi_{\mathrm{ch}}$, we refer to previous work [S2, S22].

The following parameters were used to specify $\xi_{\mathrm{ch}}$. Polar atoms contributing to $\xi_{\mathrm{ch}}$ were taken as the oxygen atoms of water and lipid phosphate moieties. The radius of the cylinder was taken as $R_{\mathrm{cyl}} = 1.2$ nm. Critically, the cylinder radius $R_{\mathrm{cyl}}$ does not impose the radius of the nucleating defect but merely ensures the locality of the defect in the membrane plane. Specifically, the use of the cylinder excludes that two laterally displaced partial defects (one from the upper and one from the lower water compartment) are misinterpreted as a continuous transmembrane defect, which would lead to undesired hysteresis problems. The cylinder was composed of $N_s = 28$ slices with a thickness of $d_s = 1$ Å. The length of the cylinder $N_s d_s$ was chosen such that $\xi_{\mathrm{ch}} \approx 0.3$ for a flat unperturbed membrane, implying that ~30% of the cylinder slices were filled by polar atoms in a flat membrane. Upon the addition of the first polar atom to a slice, the slice was considered as filled by a fraction of $\zeta = 0.75$.

The chain coordinate $\xi_{\mathrm{ch}}$ is not suitable for studying pore expansion because all pores with larger radii are projected onto $\xi_{\mathrm{ch}} = 1$. Therefore, we recently introduced a joint reaction coordinate covering pore nucleation and pore expansion (Figs. 1F/G) defined as [S26]

$$\xi_{\mathrm{p}}(\mathbf{r}) = \xi_{\mathrm{ch}}(\mathbf{r}) + H_\epsilon[\xi_{\mathrm{ch}}(\mathbf{r}) - \xi_{\mathrm{ch}}^{\mathrm{s}}] \frac{R(\mathbf{r}) - R_0}{R_0} \tag{S3}$$

Here, $H_\epsilon$ is a differentiable approximation to the Heaviside step function with switch interval $[-\epsilon, \epsilon]$. $R(\mathbf{r})$ is the radius of a fully formed transmembrane pore, and $R_0 \approx 0.42$ nm the radius of a minimal continuous defect. $\xi_{\mathrm{ch}}^{\mathrm{s}}$ indicatess the position along $\xi_{\mathrm{ch}}$ where the switch from pore nucleation to pore expansion occurs. Hence, according to Eq. S3, $\xi_p$ is equivalent to $\xi_{ch}$ for $\xi_p < \xi^s_{ch} - \epsilon$, thereby quantifying the connectivity of a nucleating transmembrane defect; for $\xi_p > \xi^s_{ch} + \epsilon$, the coordinate $\xi_p$ quantifies the radius of the open pore in units of $R_0$. Within the switch region $\xi^s_{ch} \pm \epsilon$, the $\xi_p$ gradually switches from a measure for the connectivity of the transmembrane defect to a measure for the radius of the pore (at $\xi_p \approx 0.925$ in Figs. 1F/G).

Additional parameters used for defining $\xi_p$ were chosen as follows: We used $\epsilon = 0.05$ and $\xi^s_{ch} = 0.925$. The radius $R(\mathbf{r})$ of the open pore was estimated from the number of polar atoms within a layer of thickness 1.2 nm that was positioned at the membrane center and parallel to the membrane plane, assuming a volume of 0.02996 $\mathrm{nm}^3$ per polar atom, corresponding to the volume per water molecule. For additional details were refer to Ref. S26.

#### C. Potential of mean force calculations

Initial frames for US were taken from constant-velocity pulling simulations along $\xi_{ch}$ over 50 ns for pore nucleation simulations and along $\xi_p$ over 200 ns for pore expansion. For PMF calculations along $\xi_{ch}$, US windows were spaced by 0.02 along $\xi_{ch}$ near the nucleation barrier (force constant $k = 40000\,\mathrm{kJ\,mol^{-1}}$) and spaced by 0.08 in other regions of the PMF ($k = 5000\,\mathrm{kJ\,mol^{-1}}$). For the simulations of pore expansion, the US windows were spaced by 0.08 in the early nucleation regime, by 0.03 near the nucleation barrier, and by 0.15 for $\xi_p > 1.12$. For the three regimes, force constants of 3000, 5000, and 400 $\mathrm{kJ\,mol^{-1}}$ were used. Umbrella windows for pore nucleation or for the medium-sized pore expansion system were simulated for 100 ns. Umbrella windows for the large pore expansion system were simulated for 25 ns. The PMFs were calculated using the Gromacs implementation of the weighted histogram analysis method [S27, S28]. The uncertainties of the PMFs were estimation using 50 rounds of Bayesian bootstrapping of complete histograms [S28]. For the nucleation, the initial 40 ns of each window were ignored for equilibration, whereas for the expansion, the initial 10 ns were ignored.

For PMFs covering both pore nucleation and pore expansion (Figs. 1F/G), we combined US windows from the three different DPPC simulations systems. Accordingly, windows from the small system (128 DPPC lipids) were used with US reference values $\xi_p < 0.84$ (where $\xi_p$ is equivalent to $\xi_{ch}$), windows from the medium-sized system (512 DPPC lipids) were used with reference values $0.65 < \xi_p \leq 7$, and windows from the large system (2048 DPPC lipids) were used with reference values $6 \leq \xi_p \leq 17$ (approx. 150 windows total).

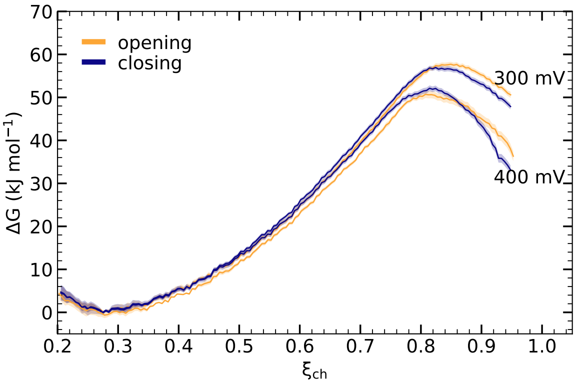


Figure S1: PMFs computed along pore opening and closing pathways exhibit only negligible hysteresis.

#### D. Additional control simulations

To validate that the PMFs are (i) converged and (ii) truly reflect the underlying free energy landscape, two additional sets of simulation were carried out.

##### *1. Absence of hysteresis between pore-opening and pore-closing pathways*

As a sensitive test for the convergence of the PMFs, we computed the PMFs in forward (pore-opening) and reverse (pore-closing) directions. Accordingly, initial frames for umbrella sampling were either taken from constant-velocity pulling simulations carried out in forward or reverse directions. Except that the constant-velocity closing simulation was carried out more slowly over 500 ns, the simulation parameters were identical. As

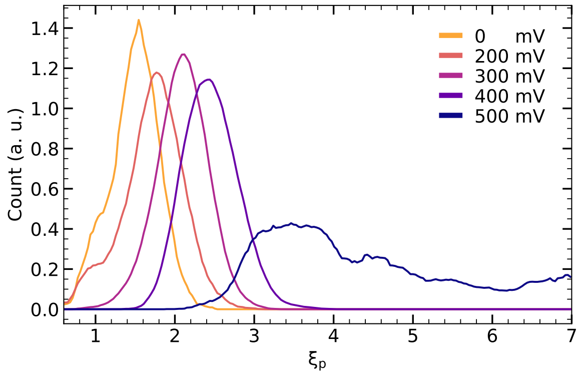


Figure S2: Distribution of $\xi_p$ in free simulations with an open pore at voltages between 0 and 500 mV, corresponding to distributions of pore radii $R$ via $R = \xi_p R_0$, where $R_0 \approx 0.42$ nm. With increasing potential the pores exhibit larger radii. At 500 mV, the pore spontaneously expanded during the simulations (dark blue).

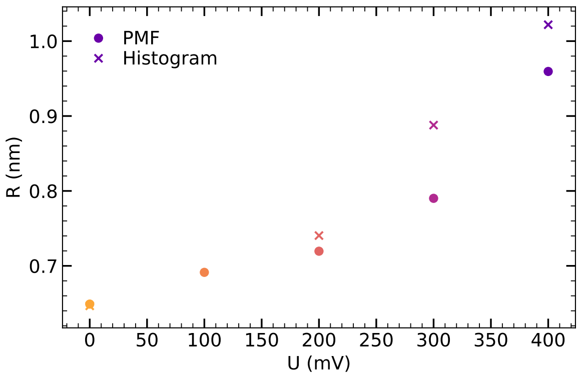


Figure S3: Comparison of the most populated $\xi_p$-positions in free simulations (crosses) as taken from the histograms in Fig. S2 with the positions of free energy minima (dots) taken from the PMFs in Fig. 1F. Color code according to Fig. S2.

shown in Fig. S1, the PMFs in forward and reverse direction favorably agree, suggesting that the PMFs are converged and that no barriers orthogonal to the RC are integrated out.

#### 2. *Validation of PMFs by free simulations*

To test whether the PMFs (Fig. 1F) reflect the true free energy landscape, we carried out free simulations at 0, 200, 300, 400, or 500 mV, starting form a conformation with an open pore. Five simulations per voltage were carried for 600 ns each. We extracted the position along the $\xi_p$ coordinate after omitting the first 30ns using frames every 250 ps and obtained the histograms along $\xi_p$ using frames over 600 ns or until the pore irreversibly closed or expanded (Fig. S2). Evidently, the average $\xi_p$ (hence the average pore radius) increased with increasing pore potential, as observed in previous simulations of DOPC/cholesterol membranes [S29].

Figure S3 presents the positions of the maxima of the histograms versus potential $U$ for voltages between 0 and 400 mV as colored crosses, compared with the positions of the local minima of the PMFs in Fig. 1F. Considering that the minina of the PMFs are shallow and, hence, their position subject to some uncertainty, the agreement between the free simulations and the PMFs is reasonable.

Furthermore, the five simulations at 0 mV closed spontaneously at simulation times between 40 ns and 140 ns, while the five simulations at 200 mV closed spontaneously at simulation times between 80 ns and 550 ns, in line with the highly shallow free energy minima in these PMFs (Fig. 1F, yellow and light red). The pores in simulations with 300 mV or 400 mV were stable over the total simulation time of 3000 ns, as expected from the pronounced free energy minima (Fig. 1F, light and dark purple). In contrast, the pore in the simulation with 500 mV rapidly expanded leading to membrane collapse within at most 500 ns, in agreement with the decreasing PMF along $\xi_p$ (Fig. 1F, dark blue). Hence, the free simulations confirm the PMFs in terms of the position and depth of the local free energy minimum of open pores.


[S1] T. Kotnik, L. Rems, M. Tarek, and D. Miklavčič, Annu. Rev. Biophys. **48**, 63 (2019).
[S2] J. S. Hub and N. Awasthi, J. Chem. Theory Comput. **13**, 2352 (2017).
[S3] K. C. Melikov, V. A. Frolov, A. Shcherbakov, A. V. Samsonov, Y. A. Chizmadzhev, and L. V. Chernomordik, Biophys. J. **80**, 1829 (2001).
[S4] C. J. Knight and J. S. Hub, Bioinformatics **31**, 2897 (2015).
[S5] O. Berger, O. Edholm, and F. Jähnig, Biophys. J. **72**, 2002 (1997).
[S6] H. J. C. Berendsen, J. P. M. Postma, W. F. van Gunsteren, and J. Hermans, in *Intermolecular Forces*, edited by B. Pullman (D. Reidel Publishing Company, Dordrecht, 1981), pp. 331–342.
[S7] M. J. Abraham, T. Murtola, R. Schulz, S. Páll, J. C. Smith, B. Hess, and E. Lindahl, SoftwareX **1**, 19 (2015).
[S8] G. Bussi, D. Donadio, and M. Parrinello, J. Chem. Phys. **126**, 014101 (2007).
[S9] M. Bernetti and G. Bussi, The Journal of Chemical Physics **153**, 114107 (2020).
[S10] T. Darden, D. York, and L. Pedersen, J. Chem. Phys. **98**, 10089 (1993).
[S11] U. Essmann, L. Perera, M. L. Berkowitz, T. Darden, H. Lee, and L. G. Pedersen, J. Chem. Phys. **103**, 8577 (1995).
[S12] S. Miyamoto and P. A. Kollman, J. Comp. Chem. **13**, 952 (1992).
[S13] B. Hess, J. Chem. Theory Comput. **4**, 116 (2008).
[S14] B. Roux, Biophysical Journal **95**, 4205 (2008).
[S15] J. Melcr, D. Bonhenry, Š. Timr, and P. Jungwirth, Journal of Chemical Theory and Computation **12**, 2418 (2016).
[S16] G. Kasparyan and J. S. Hub, bioRxiv (2023), URL https://doi.org/10.1101/2023.01.13.523896.
[S17] J. B. Klauda, R. M. Venable, J. A. Freites, J. W. O'Connor, D. J. Tobias, C. Mondragon-Ramirez, I. Vorobyov, A. D. MacKerell, Jr, and R. W. Pastor, J Phys Chem B **114**, 7830 (2010).
[S18] R. Pastor and A. MacKerell Jr, J. Phys. Chem. Lett. **2**, 1526 (2011).
[S19] C. W. Hopkins, S. Le Grand, R. C. Walker, and A. E. Roitberg, J. Chem. Theory Comput. **11**, 1864 (2015).
[S20] W. L. Jorgensen, J. Chandrasekhar, J. D. Madura, R. W. Impey, and M. L. Klein, JCP **79**, 926 (1983).
[S21] G. M. Torrie and J. P. Valleau, Chem. Phys. Lett. **28**, 578 (1974).
[S22] N. Awasthi and J. S. Hub, in *Biomembrane Simulations. Computational Studies of Biological Membranes* (CRC Press, Taylor & Francis Group, 2019).

[S23] N. Awasthi, W. Kopec, N. Wilkosz, D. Jamróz, J. S. Hub, M. Zatorska, R. Petka, M. Nowakowska, and M. Kepczynski, ACS Biomater. Sci. Eng. **5**, 780 (2019).
[S24] G. Kasparyan, C. Poojari, T. Róg, and J. S. Hub, J. Phys. Chem. B **124**, 8811 (2020).
[S25] S. F. Verbeek, N. Awasthi, N. K. Teiwes, I. Mey, J. S. Hub, and A. Janshoff, Eur Biophys J **50**, 127 (2021).
[S26] J. S. Hub, J. Chem. Theory Comput. **17**, 1229 (2021).
[S27] S. Kumar, D. Bouzida, R. H. Swendsen, P. A. Kollman, and J. M. Rosenberg, J. Comput. Chem. **13**, 1011 (1992).
[S28] J. S. Hub, B. L. de Groot, and D. van der Spoel, J. Chem. Theory Comput. **6**, 3713 (2010).
[S29] M. L. Fernández, M. Risk, R. Reigada, and P. T. Vernier, Biochemical and Biophysical Research Communications **423**, 325 (2012).
